# Whole-Genome Sequence of African Swine Fever Virus isolate from India provides insights into diversity and evolution

**DOI:** 10.1101/2022.01.31.478458

**Authors:** Sonalika Mahajan, Juwar Doley, Saravanan Subramaniam, Ajay K. Yadav, Seema Rani Pegu, Narayana H. Mohan, Vishal Chander, Karikalan Mathesh, Sukdeb Nandi, Karam Pal Singh, Vivek Kumar Gupta, Gaurav Kumar Sharma

## Abstract

African swine fever (ASF) was first reported in 1921, and since then has posed a major threat to the world pig industry and still remains a major challenge as there is no vaccine or therapy available. In May 2020, the first incidence of ASF was recorded in India, followed by a number of outbreaks in the north eastern part of India. In this study, we report the first whole genome of an Indian isolate of ASF virus (ASF/IND/20/CAD/543) using next generation sequencing and compared with the other ASFV complete genome. On phylogenetic analysis, the virus was assigned to genotype II on the basis of p72 genotyping. However, the whole genome based phylogeny distinguished it from other genotype II isolates of clade 1.1.1. This study adds to our understanding of ASFV’s genetic diversity and molecular evolution.

## 1. Introduction

African swine fever (ASF) is a World Organization for Animal Health (OIE) notifiable disease of domestic pigs and wild boars that causes high mortality, approaching up to 100% [1]. ASF is caused by ASF virus (ASFV), a large, multi-enveloped, highly complex virus belonging to the genus *Asfivirus* of the family *Asfarviridae* [2]. The genome of ASFV is composed of a linear, double-stranded DNA molecule with a size ranging between ∼170-190 kb and encodes more than 150 open reading frames (ORFs) [3]. The ASF, first described a century ago [4], remains as a challenge to control due to difficulty in development of vaccines and therapeutics [5]. The control of ASF relies on rapid diagnosis, animal stamping out, and restriction of trade and animal movements, thus leading to severe economic consequences for affected countries [6].

ASF was first reported from Kenya in 1921 and remained confined to the African continent till 1957, when it was introduced into Portugal, Europe and Cuba, Brazil, the Dominican Republic and Haiti [7]. In 2007, it was introduced into the Georgian Republic and spread to other Caucasian countries, Russia, Ukraine and Belarus in the following years [8-9]. The recent introduction of ASFV in China (2018), Myanmar (2019), and India in the year 2020 has serious economic implications and poses a major threat to the to the pork industry around the globe [10]. The North-Eastern (NE) region of India rears 47% of the Indian pig population, with an annual swine business of around US$ 0.94 billion. It has been estimated that about 54,150 pigs have died in India up to July 2021 leading to the direct losses to a tune of INR 2.76 billion [10].

In 2020, the first laboratory confirmation of ASF from India was published, and the isolates were assigned to post-2007-p72-genotype II [11] on the basis of partial characterisation of ASF virus isolates. Until now 24 genotypes of ASFV has been described globally based on the p72 gene [3, 12, 13]. However, no full genome characterization of the ASFV isolates circulating in India has been reported yet. In the present study, complete coding region of the ASFV strain (ASF/IND/20/CAD/543) obtained directly from a field sample was determined using next generation sequencing. A comprehensive genomic comparison of ASF/IND/20/CAD/543 with other ASFV whole genome from across the world shows its distinctiveness and provides insights into the evolution of ASFV and its transboundary spread.

## 2. Materials and Methods

Samples were collected from 12 pigs died of acute ASF from Dhemaji district of Assam, India in February 2021 during an outbreak and spleen tissues were collected during post-mortem examination. Nucleic acid extraction, followed by World Organization for Animal Health recommended qPCR confirmed the presence of ASFV in all samples.

### 2.1 Screening of samples for ASFV

Viral nucleic acid was extracted from the tissue homogenates using QIAamp DNA mini kit (Qiagen, Germany) as per the manufacturer’s protocol. The samples were tested for ASFV using OIE recommended protocol [14]. In brief, a 20µl reaction was assembled consisting of mastermix, probes (reference) and amplification was carried out in ABI 7500 Real-Time PCR system (Applied Biosystems, USA) with following thermal profile: 1 cycle at 95°C for 5 min, followed by 45 cycles at 95°Cfor 15 s and 58°C for 1 min.

### 2.2. Genome Sequencing

Amongst the samples, one representative isolate (ASF/IND/20/CAD/543) was selected for next generation sequencing. The genome sequencing was generated at a commercial facility (Eurofins India). Briefly, nucleic acid was extracted from the tissue sample using QIAamp DNA extraction kit (Qiagen, Germany) as per the recommended protocol. Sequencing library was prepared with TruSeq library preparation kit (Illumina, USA) followed by sequencing in a NextSeq 500 system (Illumina, USA) to generate 2 × 150bp paired reads. The sequenced raw data was processed to obtain high quality clean reads using Trimmomatic v0.38 [15] to remove adapter sequences, ambiguous reads (reads with unknown nucleotides “N” larger than 5%), and low-quality sequences (reads with more than 10% quality threshold (QV) < 20 phred score). The high quality reads of the samples were aligned to the reference ASF genome of strain BAV17 available in the NCBI database (NC_001659.2) [16] followed by annotation and variant calling. Annotation was performed by using Genome annotation transfer utility (GATU) software [17], The high quality reads of the samples were aligned to the reference sequence using Burrows–Wheeler Algorithm (software version 0.7.17) [18]. The sequence data is available in BioSample database (Accession numbers; XXXXXXX). Consensus sequence of 170,095 bp was extracted using mpileup utility of Samtools v 0.1.18 [19].

### 2.3 Phylogenetic analysis

A total of 35 genome sequences representing viral strains from different endemic countries were obtained from GenBank and the details are listed (Table 1). The pair-wise sequence alignment was performed using MAFFT [20] and nucleotide identity matrix was created using BioEdit 7.2.5 [21]. The phylogenetic analysis was performed on the basis of whole genome and three different genes namely B646L, KP177R and A238L using maximum likelihood model available in MEGA X with 1000 bootstrap replications [22]. The data set was best fitted by the nucleotide substitution model HKY+G, which was used to infer phylogenetic relationships.

**Table 1.**
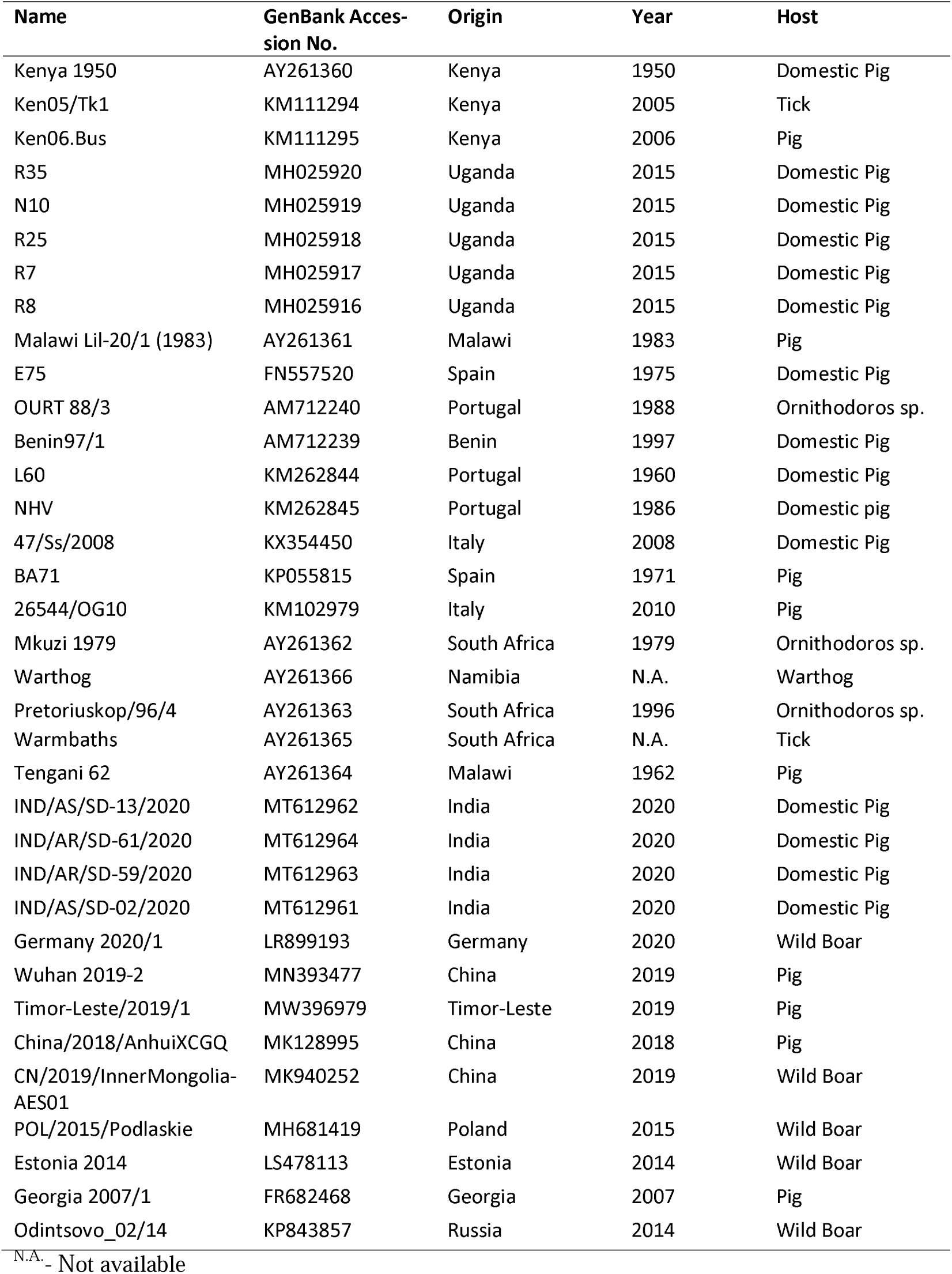
ASFV sequences used in this study

| Name | GenBank Accession No. | Origin | Year | Host |
| --- | --- | --- | --- | --- |
| Kenya 1950 | AY261360 | Kenya | 1950 | Domestic Pig |
| Ken05/Tk1 | KM111294 | Kenya | 2005 | Tick |
| Ken06.Bus | KM111295 | Kenya | 2006 | Pig |
| R35 | MH025920 | Uganda | 2015 | Domestic Pig |
| N10 | MH025919 | Uganda | 2015 | Domestic Pig |
| R25 | MH025918 | Uganda | 2015 | Domestic Pig |
| R7 | MH025917 | Uganda | 2015 | Domestic Pig |
| R8 | MH025916 | Uganda | 2015 | Domestic Pig |
| Malawi Lil-20/1 (1983) | AY261361 | Malawi | 1983 | Pig |
| E75 | FN557520 | Spain | 1975 | Domestic Pig |
| OURT 88/3 | AM712240 | Portugal | 1988 | Ornithodoros sp. |
| Benin97/1 | AM712239 | Benin | 1997 | Domestic Pig |
| L60 | KM262844 | Portugal | 1960 | Domestic Pig |
| NHV | KM262845 | Portugal | 1986 | Domestic pig |
| 47/Ss/2008 | KX354450 | Italy | 2008 | Domestic Pig |
| BA71 | KP055815 | Spain | 1971 | Pig |
| 26544/OG10 | KM102979 | Italy | 2010 | Pig |
| Mkuzi 1979 | AY261362 | South Africa | 1979 | Ornithodoros sp. |
| Warthog | AY261366 | Namibia | N.A. | Warthog |
| Pretoriuskop/96/4 | AY261363 | South Africa | 1996 | Ornithodoros sp. |
| Warmbaths | AY261365 | South Africa | N.A. | Tick |
| Tengani 62 | AY261364 | Malawi | 1962 | Pig |
| IND/AS/SD-13/2020 | MT612962 | India | 2020 | Domestic Pig |
| IND/AR/SD-61/2020 | MT612964 | India | 2020 | Domestic Pig |
| IND/AR/SD-59/2020 | MT612963 | India | 2020 | Domestic Pig |
| IND/AS/SD-02/2020 | MT612961 | India | 2020 | Domestic Pig |
| Germany 2020/1 | LR899193 | Germany | 2020 | Wild Boar |
| Wuhan 2019-2 | MN393477 | China | 2019 | Pig |
| Timor-Leste/2019/1 | MW396979 | Timor-Leste | 2019 | Pig |
| China/2018/AnhuiXCGQ | MK128995 | China | 2018 | Pig |
| CN/2019/InnerMongolia-AES01 | MK940252 | China | 2019 | Wild Boar |
| POL/2015/Podlaskie | MH681419 | Poland | 2015 | Wild Boar |
| Estonia 2014 | LS478113 | Estonia | 2014 | Wild Boar |
| Georgia 2007/1 | FR682468 | Georgia | 2007 | Pig |
| Odintsovo_02/14 | KP843857 | Russia | 2014 | Wild Boar |

<sup>N.A.</sup> - Not available

## 3. Results and Discussion

The ASFV complete genome from the Indian isolate consisted of 170,095 nucleotides with 144 Open Reading Frames (ORFs), 2,806 SNPs and had a mean GC content of 37.6%. The genome sequence of the ASFV isolate ASF/IND/20/CAD/543 has been deposited in GenBank under the accession OK236383. The pair-wise sequence alignment performed to compare ASF/IND/20/CAD/543 with other ASFV whole genomes revealed 73.3% to 86.3% identity with other sequences (Table S1). The Indian isolates sequenced in the study belonged to Genotype II and shared 100% sequence homology with isolates collected earlier from outbreaks in the Indian states of Arunachal Pradesh and Assam between January and April 2020 [11]. Till date, only genotypes I and II have been reported outside of Africa, with genotype II being responsible for the ongoing epidemic and having the most global significance. ASF genotype II viruses resulting from the Georgia introduction have been identified in a number of European countries since 2007, recently in China and Southeast Asia [3], and most recently in India [11].

The whole genome based phylogeny indicates that this virus belongs to clade 1.1.1/beta (Fig 1 b), which is consistent with the previous reports [23-24]. Previously, based on full genome analysis [23] genotype II ASFV isolates were clustered within clade 1.1.1, which also comprised genotype I. Such discrepancies between the phylogeny based on the p72 genomic region and the entire genome was reported earlier also [24]. Unlike the ASFV genotype classification based on p72 gene, presently there is no universal system for phylogeny based on complete genome analysis. Five clades (alpha, beta, gamma, delta, and epsilon), with all genotype II ASFV belonging to the beta clade was reported [25].

**Fig 1:**
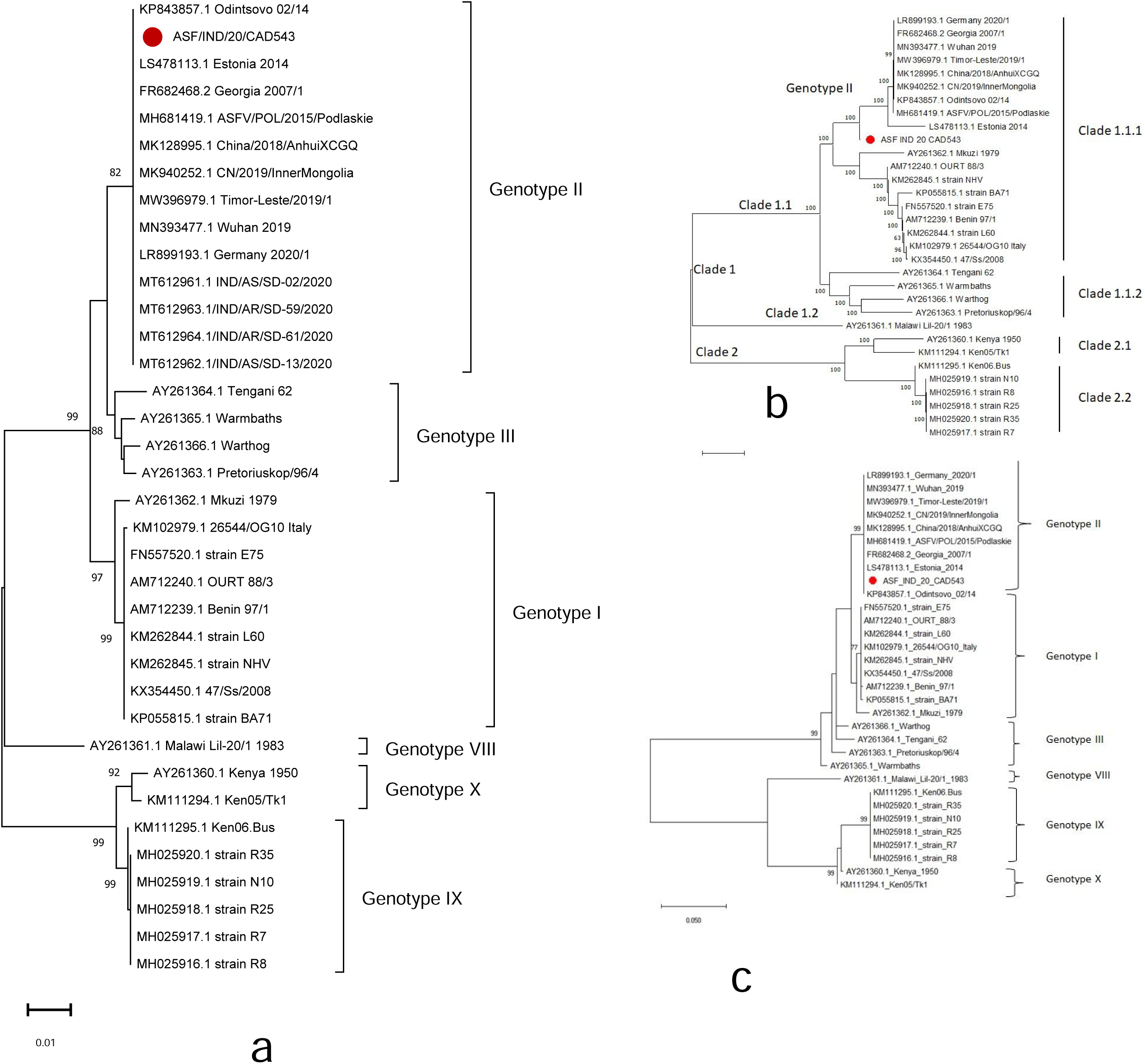
Evolutionary relationships of representative strains of African swine fever virus inferred using maximum likelihood method based on B646L(p72) gene (1a), full genome (1b) and A238L gene (1c). The uncertainty was evaluated using 1,000 bootstrap replications, and the node valuesabove 70% are shown. The isolate sequenced in this study is indicated by filled circle.

The genome size of the Indian isolate of ASF was smaller than the recently reported Tanzania/Rukwa/2017/1 isolate [3] (170095 against 183186) with less number of ORFs (144 vs 188) and more SNPs than Tanzanian or Polish or Chinese isolates. The relevance of missing ORFs is presently not known, but presumably not essential for viral replication. Furthermore, there is insertion of TRS between the I73R and I329L (Fig S1).

We also performed phylogenetic analysis based on KP177R and A238L genes. KP177R encodes p22, a structural viral protein that is early transcribed and found in the ASFV particle. Although its particular function is uncertain, p22 has been speculated to be an interaction partner of multiple host proteins [26]. The tree reconstructed using the KP177R gene exhibited a complete lack of clarity, with isolates of different genotypes clustering together and genotype II isolates collected from Estonia in 2014 constituting the basal node (data not shown). In case of A238L gene based phylogeny, the p72 genotype clustering pattern was preserved to a greater extent, with the exception of slight aberrations in genotype placement (Fig 1c). There were three primary clades in the p72 clustering pattern, with genotypes I, II, and III forming one clade, genotypes IX and X forming one clade, and genotype VIII forming one clade alone. However, in the A238L gene-based phylogeny, only two primary clades were detected, with clustered genotypes I, II, and III retained in one clade. But, the genotype VIII, which previously remained as a distinct clade in the p72-based classification, was grouped with genotypes IX and X in a single clade. The above findings suggest that the A238L gene can be used as a supplement for p72-based phylogenetic analysis. The observed differences in clustering patterns, especially with single genes, could be due to a variety of factors, including evolutionary processes that result in evolution of viral genome [27].

In the present study, we report the first whole genome based characterization of ASFV belonging to genotype II isolated from India. The isolate was assigned to genotype II and clade 1.1.1. The distinct placement of the present ASFV isolate on the basis of whole genome phylogeny indicates the genome evolution as the virus spread across the different countries with varying climate. The study reemphasizes the relevance of whole genome based characterization and classification in understanding molecular evolution of ASFV and epidemiological tracing of the disease spread.

## Acknowledgements

The authors are grateful to the Indian Council of Agricultural Research.

## Ethics Statement

The authors confirm that the ethical policies have been adhered to.

## Author Contributions

SM was involved in methodology and writing. JD, AKY, SRP, NHM, VC, KM, SN in sample procurement and data analysis. SS and GKS in conceptualization, sequencing analysis and manuscript draft. KPS and VKG in supervision and manuscript writing.

## Figure legends

Fig S1: Nucleotide sequences alignment of Indian isolate with representative strains of African swine fever virus

