## Supplementary Table for "Whole-Genome Sequence of African Swine Fever Virus isolate from India provides insights into diversity and evolution"

**Table S1**

Sequence identities matrix of the whole ASFV genomes

|  | **ASF_IND_20_CAD543** | **LR899193.1 Germany 2020/1** | **MN393477.1 Wuhan 2019** | **MW396979.1 Timor-Leste/2019/1** | **MK940252.1 CN/2019** | **MK128995.1 China/2018** | **MH681419.1 ASFV/POL/2015** | **FR682468.2 Georgia 2007** | **LS478113.1 Estonia 2014** | **KP843857.1 Odintsovo_02/14** | **FN557520.1 strain E75** | **AY261366.1 Warthog** | **AY261365.1 Warmbaths** | **AY261364.1 Tengani 62** | **AY261363.1 Pretoriuskop/96/4** | **AY261362.1 Mkuzi 1979** | **AY261361.1 Malawi Lil/1983** | **AY261360.1 Kenya 1950** | **AM712240.1 OURT 88/3** | **AM712239.1 Benin 97/1** | **KM111295.1 Ken06.Bus** | **KM111294.1 Ken05/Tk1** | **KM262844.1 strain L60** | **KM102979.1 26544/OG10 Italy** | **KM262845.1 strain NHV** | **KX354450.1 47/Ss/2008** | **KP055815.1 strain BA71** | **MH025920.1 strain R35** | **MH025919.1 strain N10** | **MH025918.1 strain R25** | **MH025917.1 strain R7** | **MH025916.1 strain R8** |
| --- | --- | --- | --- | --- | --- | --- | --- | --- | --- | --- | --- | --- | --- | --- | --- | --- | --- | --- | --- | --- | --- | --- | --- | --- | --- | --- | --- | --- | --- | --- | --- | --- |
| **ASF_IND_20_CAD543** | ID | 0.833 | 0.833 | 0.831 | 0.831 | 0.831 | 0.831 | 0.833 | 0.827 | 0.831 | 0.837 | 0.811 | 0.81 | 0.828 | 0.813 | 0.82 | 0.768 | 0.733 | 0.857 | 0.842 | 0.735 | 0.737 | 0.842 | 0.844 | 0.857 | 0.85 | 0.863 | 0.741 | 0.741 | 0.741 | 0.741 | 0.741 |
| **LR899193.1 Germany 2020/1** | 0.833 | ID | 0.997 | 0.991 | 0.993 | 0.993 | 0.993 | 0.999 | 0.942 | 0.993 | 0.901 | 0.917 | 0.933 | 0.906 | 0.918 | 0.943 | 0.848 | 0.825 | 0.855 | 0.905 | 0.828 | 0.836 | 0.907 | 0.906 | 0.856 | 0.908 | 0.886 | 0.836 | 0.835 | 0.836 | 0.836 | 0.836 |
| **MN393477.1 Wuhan 2019** | 0.833 | 0.997 | ID | 0.991 | 0.993 | 0.993 | 0.993 | 0.997 | 0.942 | 0.993 | 0.901 | 0.917 | 0.931 | 0.905 | 0.918 | 0.943 | 0.848 | 0.824 | 0.855 | 0.905 | 0.828 | 0.837 | 0.907 | 0.906 | 0.856 | 0.909 | 0.885 | 0.835 | 0.834 | 0.835 | 0.835 | 0.835 |
| **MW396979.1 Timor-Leste/2019/1** | 0.831 | 0.991 | 0.991 | ID | 0.985 | 0.985 | 0.984 | 0.991 | 0.938 | 0.984 | 0.893 | 0.91 | 0.927 | 0.902 | 0.915 | 0.94 | 0.845 | 0.821 | 0.848 | 0.898 | 0.821 | 0.83 | 0.899 | 0.901 | 0.849 | 0.906 | 0.883 | 0.832 | 0.831 | 0.832 | 0.832 | 0.832 |
| **MK940252.1 CN/2019** | 0.831 | 0.993 | 0.993 | 0.985 | ID | 0.999 | 0.999 | 0.993 | 0.936 | 0.999 | 0.906 | 0.917 | 0.93 | 0.903 | 0.915 | 0.94 | 0.848 | 0.823 | 0.86 | 0.911 | 0.833 | 0.841 | 0.913 | 0.911 | 0.861 | 0.907 | 0.884 | 0.833 | 0.832 | 0.833 | 0.833 | 0.833 |
| **MK128995.1 China/2018** | 0.831 | 0.993 | 0.993 | 0.985 | 0.999 | ID | 0.999 | 0.993 | 0.936 | 0.999 | 0.906 | 0.917 | 0.93 | 0.903 | 0.915 | 0.94 | 0.848 | 0.823 | 0.86 | 0.911 | 0.833 | 0.841 | 0.913 | 0.911 | 0.862 | 0.907 | 0.884 | 0.833 | 0.832 | 0.833 | 0.833 | 0.833 |
| **MH681419.1 ASFV/POL/2015** | 0.831 | 0.993 | 0.993 | 0.984 | 0.999 | 0.999 | ID | 0.993 | 0.936 | 0.999 | 0.906 | 0.917 | 0.93 | 0.903 | 0.915 | 0.94 | 0.848 | 0.823 | 0.86 | 0.911 | 0.833 | 0.841 | 0.913 | 0.911 | 0.861 | 0.907 | 0.884 | 0.833 | 0.832 | 0.833 | 0.833 | 0.833 |
| **FR682468.2 Georgia 2007** | 0.833 | 0.999 | 0.997 | 0.991 | 0.993 | 0.993 | 0.993 | ID | 0.942 | 0.993 | 0.901 | 0.917 | 0.933 | 0.906 | 0.918 | 0.943 | 0.848 | 0.825 | 0.855 | 0.905 | 0.828 | 0.836 | 0.907 | 0.906 | 0.856 | 0.908 | 0.886 | 0.836 | 0.835 | 0.836 | 0.836 | 0.836 |
| **LS478113.1 Estonia 2014** | 0.827 | 0.942 | 0.942 | 0.938 | 0.936 | 0.936 | 0.936 | 0.942 | ID | 0.936 | 0.904 | 0.873 | 0.888 | 0.876 | 0.881 | 0.896 | 0.824 | 0.788 | 0.825 | 0.901 | 0.795 | 0.794 | 0.901 | 0.901 | 0.825 | 0.906 | 0.889 | 0.801 | 0.8 | 0.801 | 0.801 | 0.801 |
| **KP843857.1 Odintsovo_02/14** | 0.831 | 0.993 | 0.993 | 0.984 | 0.999 | 0.999 | 0.999 | 0.993 | 0.936 | ID | 0.906 | 0.917 | 0.93 | 0.903 | 0.914 | 0.939 | 0.847 | 0.823 | 0.86 | 0.911 | 0.833 | 0.84 | 0.912 | 0.911 | 0.861 | 0.907 | 0.883 | 0.833 | 0.832 | 0.833 | 0.833 | 0.833 |
| **FN557520.1 strain E75** | 0.837 | 0.901 | 0.901 | 0.893 | 0.906 | 0.906 | 0.906 | 0.901 | 0.904 | 0.906 | ID | 0.875 | 0.888 | 0.871 | 0.886 | 0.903 | 0.827 | 0.794 | 0.896 | 0.987 | 0.815 | 0.805 | 0.99 | 0.983 | 0.897 | 0.978 | 0.949 | 0.808 | 0.808 | 0.808 | 0.808 | 0.808 |
| **AY261366.1 Warthog** | 0.811 | 0.917 | 0.917 | 0.91 | 0.917 | 0.917 | 0.917 | 0.917 | 0.873 | 0.917 | 0.875 | ID | 0.946 | 0.924 | 0.931 | 0.913 | 0.842 | 0.817 | 0.838 | 0.877 | 0.825 | 0.832 | 0.877 | 0.878 | 0.838 | 0.876 | 0.87 | 0.828 | 0.827 | 0.828 | 0.828 | 0.828 |
| **AY261365.1 Warmbaths** | 0.81 | 0.933 | 0.931 | 0.927 | 0.93 | 0.93 | 0.93 | 0.933 | 0.888 | 0.93 | 0.888 | 0.946 | ID | 0.919 | 0.937 | 0.936 | 0.849 | 0.829 | 0.843 | 0.894 | 0.826 | 0.842 | 0.894 | 0.896 | 0.843 | 0.897 | 0.876 | 0.832 | 0.832 | 0.832 | 0.832 | 0.832 |
| **AY261364.1 Tengani 62** | 0.828 | 0.906 | 0.905 | 0.902 | 0.903 | 0.903 | 0.903 | 0.906 | 0.876 | 0.903 | 0.871 | 0.924 | 0.919 | ID | 0.918 | 0.902 | 0.832 | 0.807 | 0.836 | 0.877 | 0.803 | 0.818 | 0.877 | 0.879 | 0.836 | 0.882 | 0.884 | 0.809 | 0.808 | 0.809 | 0.809 | 0.809 |
| **AY261363.1 Pretoriuskop/96/4** | 0.813 | 0.918 | 0.918 | 0.915 | 0.915 | 0.915 | 0.915 | 0.918 | 0.881 | 0.914 | 0.886 | 0.931 | 0.937 | 0.918 | ID | 0.918 | 0.844 | 0.821 | 0.843 | 0.89 | 0.82 | 0.832 | 0.892 | 0.89 | 0.844 | 0.895 | 0.875 | 0.825 | 0.824 | 0.825 | 0.825 | 0.825 |
| **AY261362.1 Mkuzi 1979** | 0.82 | 0.943 | 0.943 | 0.94 | 0.94 | 0.94 | 0.94 | 0.943 | 0.896 | 0.939 | 0.903 | 0.913 | 0.936 | 0.902 | 0.918 | ID | 0.844 | 0.827 | 0.857 | 0.907 | 0.824 | 0.837 | 0.908 | 0.912 | 0.858 | 0.917 | 0.894 | 0.834 | 0.833 | 0.834 | 0.834 | 0.834 |
| **AY261361.1 Malawi Lil/1983** | 0.768 | 0.848 | 0.848 | 0.845 | 0.848 | 0.848 | 0.848 | 0.848 | 0.824 | 0.847 | 0.827 | 0.842 | 0.849 | 0.832 | 0.844 | 0.844 | ID | 0.837 | 0.786 | 0.828 | 0.85 | 0.842 | 0.826 | 0.827 | 0.784 | 0.826 | 0.809 | 0.851 | 0.85 | 0.851 | 0.851 | 0.851 |
| **AY261360.1 Kenya 1950** | 0.733 | 0.825 | 0.824 | 0.821 | 0.823 | 0.823 | 0.823 | 0.825 | 0.788 | 0.823 | 0.794 | 0.817 | 0.829 | 0.807 | 0.821 | 0.827 | 0.837 | ID | 0.762 | 0.796 | 0.899 | 0.931 | 0.797 | 0.798 | 0.762 | 0.799 | 0.778 | 0.915 | 0.915 | 0.915 | 0.915 | 0.915 |
| **AM712240.1 OURT 88/3** | 0.857 | 0.855 | 0.855 | 0.848 | 0.86 | 0.86 | 0.86 | 0.855 | 0.825 | 0.86 | 0.896 | 0.838 | 0.843 | 0.836 | 0.843 | 0.857 | 0.786 | 0.762 | ID | 0.903 | 0.778 | 0.772 | 0.901 | 0.898 | 0.997 | 0.889 | 0.883 | 0.767 | 0.767 | 0.767 | 0.767 | 0.767 |
| **AM712239.1 Benin 97/1** | 0.842 | 0.905 | 0.905 | 0.898 | 0.911 | 0.911 | 0.911 | 0.905 | 0.901 | 0.911 | 0.987 | 0.877 | 0.894 | 0.877 | 0.89 | 0.907 | 0.828 | 0.796 | 0.903 | ID | 0.811 | 0.811 | 0.995 | 0.991 | 0.902 | 0.982 | 0.953 | 0.803 | 0.802 | 0.803 | 0.803 | 0.803 |
| **KM111295.1 Ken06.Bus** | 0.735 | 0.828 | 0.828 | 0.821 | 0.833 | 0.833 | 0.833 | 0.828 | 0.795 | 0.833 | 0.815 | 0.825 | 0.826 | 0.803 | 0.82 | 0.824 | 0.85 | 0.899 | 0.778 | 0.811 | ID | 0.91 | 0.809 | 0.807 | 0.776 | 0.8 | 0.779 | 0.975 | 0.973 | 0.975 | 0.975 | 0.975 |
| **KM111294.1 Ken05/Tk1** | 0.737 | 0.836 | 0.837 | 0.83 | 0.841 | 0.841 | 0.841 | 0.836 | 0.794 | 0.84 | 0.805 | 0.832 | 0.842 | 0.818 | 0.832 | 0.837 | 0.842 | 0.931 | 0.772 | 0.811 | 0.91 | ID | 0.81 | 0.81 | 0.771 | 0.804 | 0.782 | 0.905 | 0.904 | 0.905 | 0.905 | 0.905 |
| **KM262844.1 strain L60** | 0.842 | 0.907 | 0.907 | 0.899 | 0.913 | 0.913 | 0.913 | 0.907 | 0.901 | 0.912 | 0.99 | 0.877 | 0.894 | 0.877 | 0.892 | 0.908 | 0.826 | 0.797 | 0.901 | 0.995 | 0.809 | 0.81 | ID | 0.991 | 0.903 | 0.984 | 0.955 | 0.803 | 0.802 | 0.803 | 0.803 | 0.803 |
| **KM102979.1 26544/OG10 Italy** | 0.844 | 0.906 | 0.906 | 0.901 | 0.911 | 0.911 | 0.911 | 0.906 | 0.901 | 0.911 | 0.983 | 0.878 | 0.896 | 0.879 | 0.89 | 0.912 | 0.827 | 0.798 | 0.898 | 0.991 | 0.807 | 0.81 | 0.991 | ID | 0.897 | 0.99 | 0.958 | 0.803 | 0.802 | 0.803 | 0.803 | 0.803 |
| **KM262845.1 strain NHV** | 0.857 | 0.856 | 0.856 | 0.849 | 0.861 | 0.862 | 0.861 | 0.856 | 0.825 | 0.861 | 0.897 | 0.838 | 0.843 | 0.836 | 0.844 | 0.858 | 0.784 | 0.762 | 0.997 | 0.902 | 0.776 | 0.771 | 0.903 | 0.897 | ID | 0.891 | 0.885 | 0.768 | 0.767 | 0.768 | 0.768 | 0.768 |
| **KX354450.1 47/Ss/2008** | 0.85 | 0.908 | 0.909 | 0.906 | 0.907 | 0.907 | 0.907 | 0.908 | 0.906 | 0.907 | 0.978 | 0.876 | 0.897 | 0.882 | 0.895 | 0.917 | 0.826 | 0.799 | 0.889 | 0.982 | 0.8 | 0.804 | 0.984 | 0.99 | 0.891 | ID | 0.966 | 0.805 | 0.804 | 0.805 | 0.805 | 0.805 |
| **KP055815.1 strain BA71** | 0.863 | 0.886 | 0.885 | 0.883 | 0.884 | 0.884 | 0.884 | 0.886 | 0.889 | 0.883 | 0.949 | 0.87 | 0.876 | 0.884 | 0.875 | 0.894 | 0.809 | 0.778 | 0.883 | 0.953 | 0.779 | 0.782 | 0.955 | 0.958 | 0.885 | 0.966 | ID | 0.784 | 0.784 | 0.784 | 0.784 | 0.784 |
| **MH025920.1 strain R35** | 0.741 | 0.836 | 0.835 | 0.832 | 0.833 | 0.833 | 0.833 | 0.836 | 0.801 | 0.833 | 0.808 | 0.828 | 0.832 | 0.809 | 0.825 | 0.834 | 0.851 | 0.915 | 0.767 | 0.803 | 0.975 | 0.905 | 0.803 | 0.803 | 0.768 | 0.805 | 0.784 | ID | 0.997 | 0.999 | 0.999 | 0.999 |
| **MH025919.1 strain N10** | 0.741 | 0.835 | 0.834 | 0.831 | 0.832 | 0.832 | 0.832 | 0.835 | 0.8 | 0.832 | 0.808 | 0.827 | 0.832 | 0.808 | 0.824 | 0.833 | 0.85 | 0.915 | 0.767 | 0.802 | 0.973 | 0.904 | 0.802 | 0.802 | 0.767 | 0.804 | 0.784 | 0.997 | ID | 0.997 | 0.997 | 0.997 |
| **MH025918.1 strain R25** | 0.741 | 0.836 | 0.835 | 0.832 | 0.833 | 0.833 | 0.833 | 0.836 | 0.801 | 0.833 | 0.808 | 0.828 | 0.832 | 0.809 | 0.825 | 0.834 | 0.851 | 0.915 | 0.767 | 0.803 | 0.975 | 0.905 | 0.803 | 0.803 | 0.768 | 0.805 | 0.784 | 0.999 | 0.997 | ID | 0.999 | 0.999 |
| **MH025917.1 strain R7** | 0.741 | 0.836 | 0.835 | 0.832 | 0.833 | 0.833 | 0.833 | 0.836 | 0.801 | 0.833 | 0.808 | 0.828 | 0.832 | 0.809 | 0.825 | 0.834 | 0.851 | 0.915 | 0.767 | 0.803 | 0.975 | 0.905 | 0.803 | 0.803 | 0.768 | 0.805 | 0.784 | 0.999 | 0.997 | 0.999 | ID | 0.999 |
| **MH025916.1 strain R8** | 0.741 | 0.836 | 0.835 | 0.832 | 0.833 | 0.833 | 0.833 | 0.836 | 0.801 | 0.833 | 0.808 | 0.828 | 0.832 | 0.809 | 0.825 | 0.834 | 0.851 | 0.915 | 0.767 | 0.803 | 0.975 | 0.905 | 0.803 | 0.803 | 0.768 | 0.805 | 0.784 | 0.999 | 0.997 | 0.999 | 0.999 | ID |
